# T1K: efficient and accurate KIR and HLA genotyping with next-generation sequencing data

**DOI:** 10.1101/2022.10.26.513955

**Authors:** Li Song, Gali Bai, X. Shirley Liu, Bo Li, Heng Li

## Abstract

Killer immunoglobulin-like receptor (KIR) genes and human leukocyte antigen (HLA) genes are highly polymorphic in a population and play important roles in innate and adaptive immunity. We have developed a novel computational method T1K that can efficiently and accurately infer the KIR or HLA alleles from next-generation sequencing data. T1K is flexible and is compatible with various sequencing platforms including RNA-seq and genomic sequencing data. We applied T1K on CD8+ T cell single-cell RNA-seq data, and identified that KIR2DL4 allele expression levels were enriched in tumor-specific CD8+ T cells.

## Background

Polymorphisms in immune receptor genes diversify the immune response, which strengthens the resilience of a population to diseases. In particular, major histocompatibility complex (MHC) encoded by the highly polymorphic human leukocyte antigen (HLA) genes can present various peptides depending on the personal HLA sequences. The peptides on MHC could trigger different immune responses thus affecting the severity of a disease like SARS-CoV-2 infections [1] in a person. Besides HLA genes, the killer immunoglobulin-like receptor (KIR) gene family residing on 19q13.4 is also highly polymorphic and can modulate the activity of natural killer (NK) cells and T cells [2]. There are 17 KIRs in humans, including eight inhibitory KIRs (KIR2DL1, KIR2DL2, KIR2DL3, KIR2DL5A, KIR2DL5B, KIR3DL1, KIR3DL2 and KIR3DL3), seven activating KIRs (KIR2DS1, KIR2DS2, KIR2DS3, KIR2DS4, KIR2DS5, KIR3DS1 and KIR2DL4), two pseudogenes (KIR2DP1 and KIR3DP1). Regarding the regulation function, KIR2DL4 is special, in that it exerts the activating signal and also has the potential for inhibition [3]. While most of the KIRs interact with MHC class I molecules, the KIR ligand space could be broad, and this involves KIRs in various immune regulation mechanisms. For example, one immune evasion mechanism in several tumor types is to upregulate HHLA2 which binds with KIR3DL3 to inhibit T cell and NK cell activity [4]. In sum, the identification of the alleles, or genotyping, of these highly polymorphic genes in a person can lead to a better understanding of infectious diseases [5], vaccination [6], organ transplantation [7, 8], autoimmune diseases [9] and cancer [10, 11].

Because of the importance of these polymorphic genes, researchers created the Immuno Polymorphism Database (IPD) to curate the HLA and KIR allele sequences (IPD-IMGT/HLA, IPD-KIR respectively) [12]. IPD-IMGT/HLA is the foundation for numerous computational methods, such as seq2HLA [13], OptiType [14], PolySolver [15], HISAT-genotype [16] and arcasHLA [17], that can infer HLA alleles from RNA-seq, whole genome sequencing (WGS) or whole exome sequencing (WES) data. However, many of the HLA genotypers have hardwired the HLA information in the program or require specialized reference sequences, so they could not be directly applied to KIR genotyping. Besides, KIR genes have unique biological features that make the sequence analysis more complicated than HLA genes. First, KIR genes can be lost on a chromosome except for the four framework KIR genes (KIR2DL4, KIR3DL2, KIR3DL3 and KIR3DP1) [18], while the HLA class I (HLA-A, HLA-B, HLA-C) and HLA class II (including HLA-DPB1, HLA-DQB1, HLA-DRB1) genes are expected to be present. Second, some KIR genes are highly similar to each other, such as KIR3DL1 and KIR3DS1, and HLA genes are more distinct (Figure S1). These differences indicate that computational KIR genotyping is more challenging. In other words, if a method can resolve the KIR genotyping well, it can handle HLA genotyping. Motivated by this rationale and the biological relationship between KIRs and HLAs, we developed the flexible and user-friendly computational method T1K (The ONE genotyper for Kir and hla) to accurately genotype KIR and HLA genes of a sample from the genomic or RNA sequencing data.

KIR genes are selectively expressed in T cells including CD8+ T cells [19]. T cell is the central component in adaptive immunity, and CD8+ T cells can recognize antigens presented on MHC class I and eliminate corresponding cells such as cancer cells. Because KIR can regulate these important CD8+ T cells, it is fundamental to understand the KIR expression patterns. Therefore, we utilized public single-cell RNA-seq (scRNA-seq) data [20] of the tumor-infiltrating CD8+ T cells to check whether certain KIR alleles could be related to tumor immunity.

## Results

### Overview of the method

The principle of T1K is to find abundant alleles and genes based on the read alignments (Figure 1a). A similar strategy is adopted in methods like HISAT-genotype [16] and arcasHLA [17]. The allele sequences can be obtained from the IPD or a custom database. T1K first extracts candidate reads from the raw data FASTQ files or an alignment BAM file. T1K computes the abundance of all the input alleles simultaneously using the weighted expectation-maximization (EM) algorithm [21] to maximize the likelihood of read alignments to the reference alleles. By modeling all the alleles together, T1K can handle the reads that are mapped to multiple highly similar KIR genes. T1K reports the allele at the allele series level (3 digits for KIR and 6 digits for HLA by default), so it will sum the abundances of each allele within the same allele series. For simplicity, we will continue using the term allele for the genotyping results. T1K assumes at most two alleles per gene, and selects the pair of alleles that maximizes the total number of compatible reads in case of more than two valid alleles. T1K then applies a Poisson model to calculate the quality score for each called allele to further filter the false alleles. In addition to genotyping each sample, T1K provides post-processing methods to extend the genotyping results. These include novel single nucleotide polymorphism (SNP) detection on representative alleles and the report of single-cell level allele abundances.

**Figure 1.**
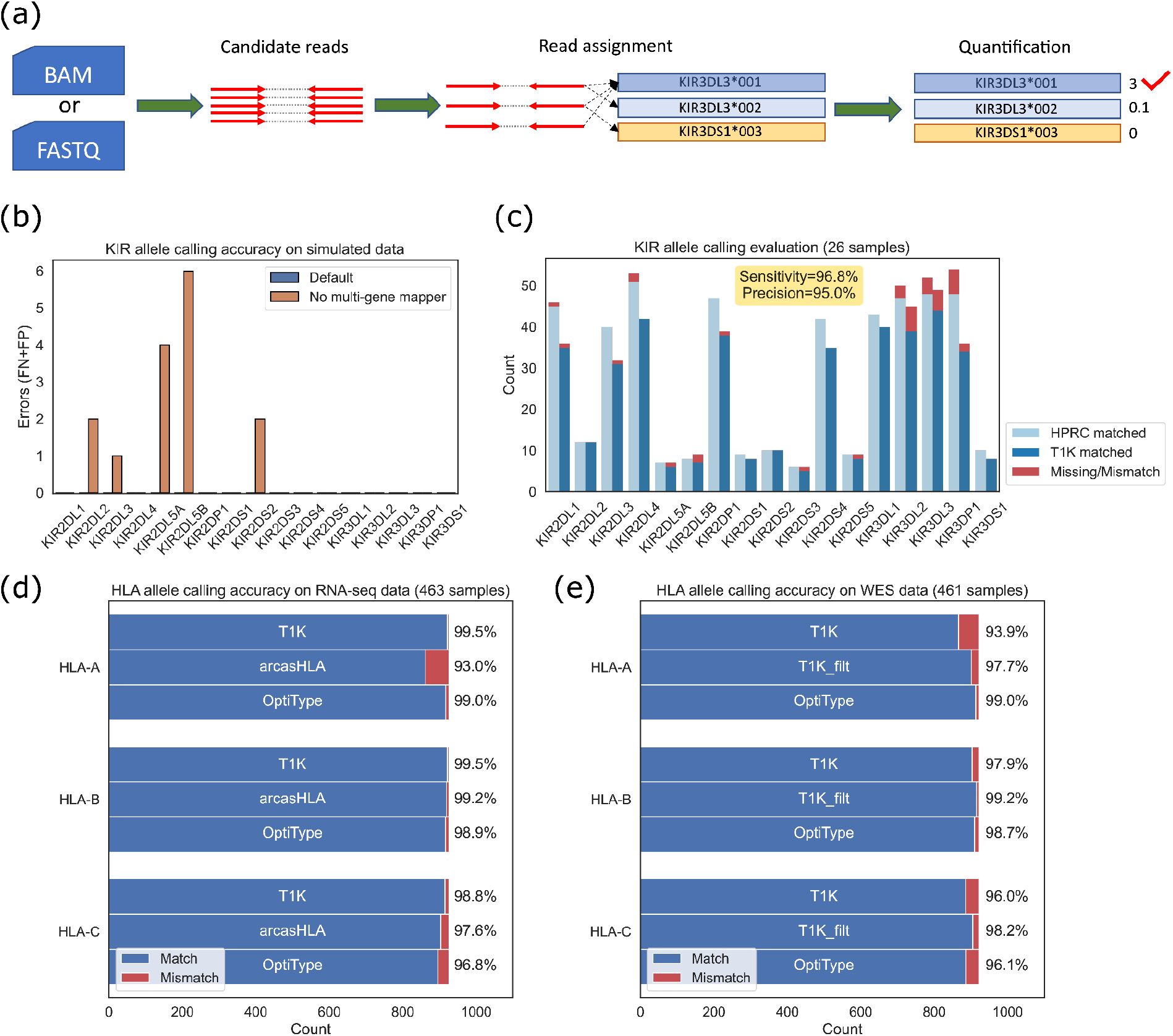
Evaluation of T1K. (a) Overview of the T1K workflow. (b) The KIR allele prediction accuracy of T1K on simulated reads from KIR mRNAs. FN: false negative, FP: false positive. (c) The KIR allele prediction accuracy of T1K on WGS. HPRC matched: the alleles inferred from the HPRC phased genome that can be found in T1K predictions. T1K matched: the alleles predicted by T1K that can be found on HPRC genome. (d) The HLA allele prediction accuracy of T1K, arcasHLA and OptiType on RNA-seq data validated with 1kPG annotation. (e) The HLA allele prediction accuracy of T1K and OptiType on WES data validated with 1 kPG annotation. T1 K_filt represents the prediction accuracy of T1K when using reduced reference sequences filtered by the OptiType database.

### Performance of KIR genotyping on simulated data

We examined the KIR genotyping accuracy of T1K with 100 simulated KIR-specific RNA-seq data. T1K made no error for the 1,642 alleles in all the simulated samples. One of the key ideas in T1K is to estimate the abundances of all the alleles together to include multiple-gene mapped reads. To test the benefit of this strategy, we removed the reads assigned to multiple genes, which is a filter adopted in arcasHLA. After the filtering, 14 samples had wrong genotyping results, supporting the importance of considering all the reads (Figure 1b). In this and following evaluations, we ignored alleles with a quality score of 0 in T1K.

To handle absent KIR genes or homozygous genes, T1K filters an allele if its abundance is lower than a user-specified fraction of the abundance of another allele for this gene (default fraction=0.15). If the threshold is set too low, then T1K will have too many candidate alleles to have robust genotyping results. If the filter threshold is too high, it will lose true alleles. We evaluated the impact of different thresholds on the allele prediction accuracy. T1K produced perfect allele predictions in a wide range of threshold settings between 0.1 and 0.25, suggesting that T1K is robust to the selection of this parameter (Figure S2).

PING [22, 23] is another pipeline for KIR genotyping. However, this pipeline reported errors on our data and its own test data, and generated “NA” for all the KIR genes in the final output folder. We have tested PING on multiple platforms (Linux, macOS; on a local machine or a server) and got “NA” on all the tests. We could not compare the results with PING in the evaluation.

### Performance of KIR genotyping on real data

We investigated T1K’s accuracy of KIR genotyping on real data. There is no public sequencing data with annotated KIR information. To conduct a comprehensive evaluation, we used 26 samples from Human Pangenome Reference Consortium (HPRC) [24, 25] (Table S2). These samples have both phased reference genomes and Illumina whole-genome sequencing short reads. To identify the KIR alleles on the phased genomes, we aligned the IPD-KIR allele genome sequences to each genome. The alignment could be used to identify KIR gene regions and to select the alleles with minimal differences in the exonic region as the ground truth (Methods). We then used T1K to predict alleles from the Illumina short reads, and compared the results against the ground truth. T1K achieved high sensitivity and identified 96.8% of alleles found on the phased reference genomes (Figure 1c). While being highly sensitive, T1K also had high precision that about 95.0% of the called alleles can be validated in the phased genomes. The phased reference genomes also provide the ground truth of novel genomics variations. We evaluated the unambiguous SNPs in the correct alleles predicted by T1K and found that 95.0% of these SNPs inferred by T1K can be validated on the phased genome (Figure S3). T1K could not resolve the ambiguous SNPs and did not try to compute the SNPs in the case of hybrid genes, so in such cases, the sensitivity was low. In this evaluation, the unambiguous SNPs of T1K constituted about 41.2% of the total novel variations found on the phased genome.

### Performance of HLA genotyping on real data

We compared T1K against other HLA genotypers to further evaluate T1K on real RNA-seq and WES data. Since arcasHLA and OptiType were reported as the most effective methods in several HLA genotyping benchmark studies [17, 26, 27], we included these two methods in our evaluations. We utilized the HLA annotation from 1000 Genome Project (1kGP) [28] as the ground truth [29] and compared the performance on samples with both RNA-seq data and WES data. Since the HLA annotation of 1kGP was at the 4-digit level, e.g. HLA-A*01:01, we converted the results from T1K and arcasHLA from 6 digits to 4 digits. Because OptiType only genotyped HLA class I genes, we focused on the analysis of the HLA-A, HLA-B and HLA-C genes. It is not likely to miss an HLA class I gene on a chromosome. Hence we regarded the single-allele (homozygous) prediction of T1K as one allele showing up twice. This is the paradigm of arcasHLA, OptiType and the 1kGP annotation. For the matching criteria, we considered each HLA gene’s alleles as a pair to avoid overcounting the matches of the homozygous case. For example, when T1K predicted the single-allele HLA-A:01:01 for a sample, and the 1kGP annotation was HLA-A*01:01; HLA-A*02:01, we counted it as one match and one mismatch. Based on these criteria, there was no need to distinguish between sensitivity and precision, which both equal the fraction of matched alleles. We first evaluated the results of T1K, arcasHLA and OptiType on the RNA-seq data (Figure 1d). T1K achieved the highest accuracy across all three genes with around 99.2% overall accuracy.

We next compared the results of T1K and OptiType on WES data. arcasHLA was excluded from this evaluation as it was incompatible with genomic sequencing data. T1K’s allele prediction accuracy was worse than OptiType, where T1K and OptiType had 96.0% and 97.9% average accuracy, respectively (Figure 1e). T1K’s accuracy was much lower than the numbers seen in the RNA-seq samples from the same donors. In RNA-seq data, a read or a read fragment can span multiple exons and can link allele-specific sequence information far apart. However, the genomics sequencing data, like WES data, could only provide local sequence information. OptiType has its own processed reference sequences with 6,889 HLA class I alleles created in 2014. Whereas T1K conducted the genotyping based on the IPD-IMGT/HLA updated in 2021 which contains 11,438 HLA class I alleles. Due to the read span limitation of WES data, we hypothesized that the prediction accuracy would deteriorate given a more complex allele reference file. To validate this assumption, we filtered the reference sequences in T1K and only kept the alleles presented in the OptiType library. The average allele prediction accuracy of T1K on the filtered database achieved 98.4% which was comparable to OptiType’s performance (Figure 1e). T1K’s ability to obtain high precision on RNA-seq data from the same donors implied that the HLA annotations in 1kGP remained valid by the standard of a more recent IPD-IMGT/HLA. Therefore, the lower accuracy on WES data before might be mostly due to that the complete database was too complicated for WES to resolve instead of annotation errors.

### Speed and memory usage

As the consequences of the comprehensiveness of IPD-IMGT/HLA and the high expression of HLA genes, HLA genotyping using RNA-seq data is the most time-consuming and memory-demanding task in our study. Therefore, we compared the computation efficiency of T1K, arcasHLA and OptiType on the ten largest RNA-seq samples that were used in the HLA genotyping evaluation (Table S1). Both T1K and OptiType started from the raw read FASTQ file, and arcasHLA used the aligned BAM file as input. All three methods could finish each sample within three hours given eight threads. Since arcasHLA extracted the candidate reads directly from the reads mapped to chromosome 6 in the BAM file, its overall running time was faster than T1K and OptiType. When comparing the speed on the genotyping step after obtaining candidate reads, T1K and arcasHLA had similar running times. While T1K and arcasHLA required less than 40G memory, Optitype consumed more than 200G memory on three of the benchmarked samples.

### Expression of KIRs in CD8+ T cells

The advancement of scRNA-seq technology enables us to explore the KIR expression patterns in CD8+ T cells computationally. To avoid the confounding effects of dropout events from low-coverage platforms like 10x Genomics scRNA-seq data [30], we chose SMART-seq [31] data for robust genotyping and abundance estimation. We analyzed 32 CD8+ T cells with SMART-seq scRNA-seq data to investigate the KIR allele expression patterns in T cells [20]. These cells were from multiple donors including cancer patients, so we first grouped the cells belonging to the same person by matching the HLA class I genes based on T1K’s results. After that, we ran T1K again on the cells from the same donor for KIR genotyping and obtained the allele abundance. To compare the KIR allele expression pattern, we normalized the abundances to expression fractions within each cell (example in Figure 2a, full results in Figure S4). We observed that not all KIR alleles or KIR genes were expressed in each cell. This was in line with the previous study that KIR genes were selectively expressed in CD8+ T cells [19] (Figure 2b). In particular, most cells expressed less than seven KIR alleles. KIR2DL4 and KIR3DL2 genes were detected in the most number of cells (27 and 21 respectively), and they were the only KIR genes that had both alleles expressed in a cell (Figure 2c). These two KIR genes were two of the four framework KIR genes, while another framework KIR gene KIR3DL3 was much less expressed. Furthermore, the remaining framework KIR gene KIR3DP1 was not expressed by any cells, implying that the pseudogene might not be functional in CD8+ T cells.

**Figure 2.**
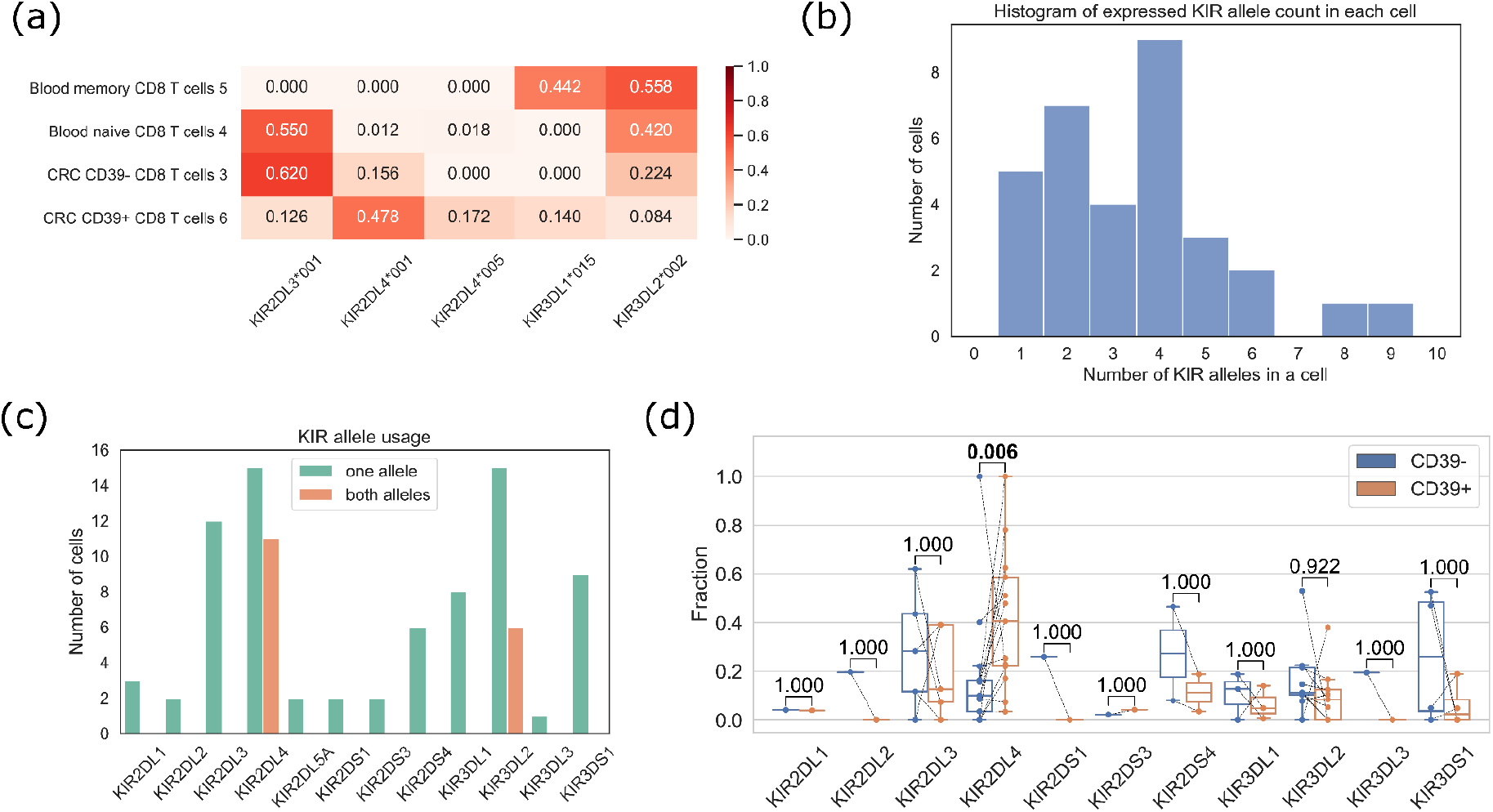
KIR alleles in CD8+ T cells. (a) KIR allele expression fractions in four cells from a colorectal cancer (CRC) patient (b) The number of expressed KIR alleles in a cell. For example, seven cells expressed two KIR alleles. (c) The number of cells that express the KIR gene, splitting by the cases of single-allele expression or both-allele expression. Only detected KIR genes are displayed. (d) Comparison of the KIR allele fractions between CD39-CD8+ T cells and CD39+ CD8+ T cells. Each line connects a KIR allele in the CD39-cell and the CD39+ T cell from the same patient. The p-values are computed with Wilcoxon signed-rank test, and have been adjusted by the Benjamin-Hochberg procedure.

We next investigated the KIR allele expression pattern in different phenotypes of the tumor-infiltrating CD8+ T cells, where CD39+ T cells are tumor-specific T cells and CD39-T cells are bystanders [20]. Since a donor contributed at most one CD39-cell and one CD39+ cell in this data, we considered each KIR allele expression fraction of a patient as a pair (CD39-versus CD39+) and ignored the patients without paired cells. When comparing the allele expression fractions, KIR2DL4 alleles were significantly enriched in the tumor-infiltrating CD39+ T cells (two-sided Wilcoxon signed-rank test, raw p-value=5.5e-4, adjusted p-value=0.006, Figure 2d). The enrichment of KIR2DL4 remained significant when applying the unpaired two-sided Mann-Whitney U test (p-value=0.013). Since KIR2DL4 could initiate the cell activating pathway, the observation of a higher KIR2DL4 fraction in tumor-associate T cells suggested it might be beneficial in CD8+ T cell anti-tumor immunity.

## Discussion

We have conducted comprehensive evaluations to show that T1K is a highly accurate genotyping method. For example, we utilized phased genomes from HPRC to validate the allele predictions. These approaches are computation-based, and they might be less reliable than experimental techniques, such as using PCR with allele-specific primers to examine the alleles. To increase our confidence in T1K, we investigated the HLA genotyping by evaluating with 1kGP annotations which were experimentally validated extensively in the original study. As KIR genes have their own unique features, experimental validation might still be needed for KIR genotyping, but this is beyond the scope of the current study.

While HLA and KIR allele sequences are well curated by IPD, there are other polymorphic genes. Some polymorphic genes like CYP2D6 relating to drug metabolism [32] are cataloged by the PharmVar database [33]. T1K provides flexible and user-friendly modules to create custom references to genotype these non-IPD curated genes. In addition, KIR genes are in the leukocyte receptor complex (LRC), and LRC contains other immune receptor gene families such as LILR and LAIR. It has been shown that a subset of the regulatory receptor genes LILR on neutrophils are genetically diverse in the population [34]. To analyze the polymorphisms in the gene without a curated database, one could retrieve the allele sequences from resources like the HPRC genomes and create a custom database for T1K to genotype new samples.

There could be structural polymorphisms in KIR loci that are too complex to process with the current implementation, such as duplicated or hybrid KIR genes [35]. T1K could not align the reads to hybrid KIR genes due to the high divergence to the reference sequences. This might decrease the allele prediction accuracy and the power to detect variations. Future works like graph representations of the KIR gene sequences at the exon level could resolve these complex variations.

In addition to demonstrating T1K’s functionality, we explored the KIR expression patterns in CD8+ T cells. We observed that KIR2DL4 was enriched in tumor-specific CD39+ CD8+ T cells suggesting that KIR2DL4 could be a therapeutic target to promote the activity of CD8+ T cells in the tumor environment. We did not explore KIR genes on 10x Genomics scRNA-seq data due to the concern over dropouts, but the analysis might be feasible and reliable at the cell cluster level. With the expanding knowledge of KIRs’ function, we expect the genotypes inferred from T1K could be a valuable resource for biologists to study and find appropriate cancer treatment strategies in the future.

## Conclusion

We have implemented the novel computational method T1K that can genotype KIR genes or HLA genes from various sequencing platforms including RNA-seq, WES and WGS data. We showed that KIR2DL4 alleles were more expressed in tumor-specific CD8+ T cells than the bystander tumor-infiltrating CD8+ T cells. T1K is free and open source at https://github.com/mourisl/T1K, and its versatile framework will contribute to KIR, HLA and other polymorphic gene studies in the future.

## Methods

### Sequencing data

We generated 100 simulated samples using Mason [36]. Each sample was generated with options “illumina -i -s 17 -sq -mp -n 100 -ll 500 -hs 0 -hi 0 -pi 0 -pd 0 -pmmb 0.0005 -pmm 0.001 -pmme 0.003 -nN -N read_count”. Each sample selected 6 to 9 random KIR genes and two random alleles (allowing the same) for each KIR gene. The parameter “read_count” in the mason command was set to have 50 read pairs for each allele on average. The read length is 100bp.

For the real data set in the KIR genotyping evaluation, we used 26 samples (Table S1) from HPRC with both haplotype-resolved reference genomes and Illumina WGS paired-end short read of 150bp read length. For the HLA genotyping comparison, we downloaded 463 samples from 1kGP that have annotated HLA genotypes, RNA-seq and WES-seq data (Table S2). We used STAR [37] with default parameters to align the RNA-seq data to the human reference genome hg38. Two of the WES samples (HG00104 and NA18487) had damaged FASTQ files, so we excluded them from the WES genotyping analysis.

### Reference sequences

IPD curates a comprehensive set of allele sequences for both HLA and KIR genes, so T1K builds the reference sequences based upon it as many other methods. In this work, T1K used IPD-KIR v2.10.0 for KIR genotyping and IPD-IMGT/HLA v3.44.0 for HLA genotyping. OptiType has its own processed HLA reference sequence in the package, and we run arcasHLA with IPD-IMGT/HLA v3.44.0. In this study, we used T1K v1.0.0, arcasHLA v0.2.5 and OptiType v1.3.5.

The T1K reference database is prepared differently depending on whether the input sequences contain introns. For RNA-seq data, T1K concatenates the exons parsed from the EMBL-ENA formatted “dat” file for each allele. For WGS or WES, we added the 250bp from each flanking intron on two sides of every exon. If two exons’ flanking introns overlap, we will merge the intervals. For the long introns, we add one character “N” at the boundary to indicate the gap in the intronic region. For all data types, we padded the sequences with 50bp sequences from 3’ and 5’ UTRs to give more anchors for the read alignment. The “dat” file also annotates whether a sequence is partial, and T1K will ignore those partial sequences.

When the UTR is exonized, such as HLA-A*04:09N, the allele contains sequences that are missing in other alleles. As a result, reads from the UTR region are more likely to be mapped to the UTR-exonized alleles and cause inflation of abundance estimation. To alleviate the mapping bias, T1K trims the last exon of an allele if the allele is longer than the common allele length of the gene. The trimming is only applied to the last exon, and the trimmed last exon length equals this gene’s most frequent last exon length. In this way, T1K will keep the sequence if there are insertions in the middle of the allele.

### Candidate read

The first step of T1K is to extract the reads from the interested genes provided in the reference sequences, such as KIR and HLA genes. For user convenience, T1K is compatible with both aligned BAM input and FASTQ input, and the choice of input format does not affect the genotyping outcome in our tests. For reads that are aligned to alternate contigs, unmapped or in FASTQ file, the extraction algorithm is similar overlap detection algorithm in TRUST4 [38] by checking the colinear seeds hit. Suppose the total length of the reference sequences is L, the seed length used in this step is log_4_L + 1. For example, in the RNA reference sequence of HLA genotyping, L is about 16 million, so seed length equals 13. As a result, a k-mer in a read hits about once to the reference sequence in expectation. Since reference sequences are highly redundant, the seed length overestimates the hit probability and will not incur much computation overhead. A read will be a candidate if the chain covers 20% of the read length. When given the BAM input, most of the reads mapped outside the genotyped genes can be directly ignored, so T1K was about 30% faster than the FASTQ input.

### Read assignment

To conduct abundance estimation, T1K assigns each read, or a read fragment, to its best-aligned alleles. The best alignments are the ones with the most matched nucleotides between the reference sequence and a read. The reference sequences are highly redundant, and a read can be aligned to thousands of alleles. This becomes the computation bottleneck in RNA-seq analysis, where HLA genes can express millions of reads. We notice that the ultra-high coverage will create a large number of identical reads. Therefore, T1K will first sort the reads by the nucleotide sequence, and then only conduct the alignment for the duplicates once. For paired-end data, T1K conducts the same procedure by regarding each read fragment as two independent read ends, and then selects the two compatible read-end alignments resulting in the optimal read pair alignment score.

T1K further supplements the read alignment by strategically incorporating assignments with lower scores. In the reference sequences for genomic sequencing data, we only include part of the introns flanking the exons and mark the remaining parts as gaps. This strategy could create alignment bias near the boundary. For example, if allele A has a deletion at position x before the gap, allele A’s intron will contain the first nucleotide of the corresponding gap in other alleles. As a result, read aligned after position x and partially overlapping with the gap will have one more base matched with allele A than other alleles. To reduce this bias, T1K considers the alignment part extending in the gap as all matches. Furthermore, the IPD-KIR is much less comprehensive than IPD-IMGT/HLA, and the inadequate information on intron sequences could lead to false negatives. Therefore, we add the option to incorporate the read assignment allowing more mismatch in the non-exonic region than the best alignment. In our evaluation with HPRC phased genomes, this strategy improves the accuracy by 2.6%.

To avoid low-quality reads and false positive read assignments, T1K filters the assignments with a low alignment identity. In the case of KIR genotyping, we require at least 80% of matched bases. For HLA genotyping, we have tested several cutoffs and found 97% gave a good performance, where 97% was also the threshold proposed in OptiType. For WGS data, since reads can come from the whole genome and are more likely to introduce false positives, we need to use a more stringent alignment similarity threshold at 90%. We have made these settings in the “--preset” option of T1K.

### Abundance estimation

T1K estimates the abundances of all the alleles by maximizing the likelihood of read assignments. The log-likelihood function is 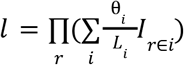, where θ_*i*_ is the proportion or probability for allele i, *L_i_* is the length of allele i, and *I_r∈i_* is 1 if allele i is one of read r’s alignment target and 0 otherwise. We apply the EM algorithm to find the solution by introducing the latent variable *z_r,i_* indicating read r is from allele i. So the conditional expectation of the log-likelihood function can be written as 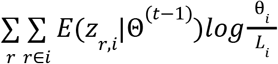 in E-step at iteration t, where 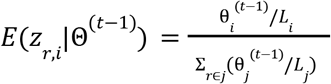. In M-step, we compute the updated abundances that maximize the conditional expectation, which is 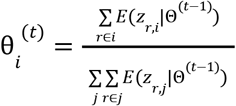. In the above equations, we use i, j to index alleles and r to index the read. The initial abundance for each allele is proportional to the allele series frequency in the database. For example, HLA-A*01:01:01:01 is in the HLA-A*01:01:01 series and the size of this series is 91 in the reference sequences for RNA-seq data, so its initial abundance is 91 times higher than the singleton HLA-A*01:82 allele. The intuition is that the large allele series suggests that it is better studied and could be more common in the population. The iterations terminate when the update changes the Θ by less than 1e-5 in total or the number of iterations exceeds 1000. Finally, the abundance for allele i is 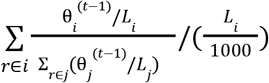 which is the definition of the fragment per thousand bases (FPK).

T1K implements two methods to expedite the execution of the EM algorithm. First, we group the reads assigned to the same set of alleles together. Therefore, the EM algorithm is weighted. Second, we adopt the SQUAREM algorithm [39] which has a faster convergence rate than the vanilla EM algorithm. To impose sparsity, we implement the heuristics that removes the low abundant alleles every 10 iterations based on the abundances estimated in that iteration. This is similar to the sparsity method in arcasHLA.

### Allele selection

The EM algorithm calculates the abundance for each allele with all digits. Because T1K does not focus on the variations in introns, it reports the alleles at a higher level with fewer digits. For example, T1K reports KIR and HLA alleles at 3-digit and 6-digit levels respectively. The abundance for each higher-level allele series is the summation of the allele abundances in the same series. For simplicity, we still indicate the allele series as an allele. T1K selects the allele with the highest abundance as the dominant allele, and filters other alleles with abundances less than a user-specified fraction (default 15%) of the dominant allele.

While the abundance filter can remove the noise allele in most cases, there could still be more than two alleles passing the abundance filter. T1K refines the selection by picking a pair of alleles that can improve the number of valid read assignments. Starting from the two most abundant alleles for each gene, T1K changes one of the alleles to find the allele that increases the read assignments the most. T1K repeats this procedure based on the pairs of alleles selected in the previous iteration until the number of explained read assignments could not be improved. This strategy favors the two most abundant alleles and is faster than enumerating all the allele pairs in most cases.

### Genotyping quality score

T1K also scores each called allele to represent the confidence. The principle is to conduct the statistical test that compares the computed abundance with the noise abundance. The noise abundance for one allele A of gene *G_i_* is calculated as Σ_*j*≠*i*_, α*Abund*(*G_j_*)*sim*(*G_i_, G_j_*) + β*Abund*(*B*), where *Abund*(*G_j_*) is the abundance of gene j; *sim*(*G_i_, G_j_*) is the sequence similarity between genes *G_i_* and *G_j_*; *Abund*(*B*) is abundance for the other allele of gene *G_i_* if it is heterozygous; α, β are user-defined constant to control the magnitude of noise. We model the abundance as Poisson distribution, and compute the p-value by testing the observed abundance under the hypothesis of noise Poisson distribution. The quality score is reported as −log_10_(p-value). Similar to the MAPQ in read alignment software like BWA-MEM [40], we set 60 as the upper limit of the quality score.

### SNP detection

Even though T1K reports the genotyping results at the allele series level, it keeps track of the actual alleles in the process and reports one allele per series as the representative. This information can be used to identify SNPs that are missing from the reference database. For SNP detection, we realign the reads to the representative alleles. We process a position i of allele A with a high mismatch rate, that is the number of alignments supporting an alternative nucleotide is at least half of the number for the reference nucleotide. Due to allele similarities, these reads could still be equally good assigned to other alleles. Therefore, T1K utilizes the multiple-mapped reads to incorporate other alleles and piles the allele sequences around position i. This procedure can be thought of as read-alignment-guided multiple sequence alignment. The SNP could happen to any of the incorporated alleles. To select the alleles containing SNPs, T1K follows the law of parsimony by introducing the least number of SNPs to explain the most read assignments (Figure S5). In more detail, T1K tries all the nucleotide combinations at position i across the piled alleles to check how many reads can have matched bases at position i in any of the alleles. T1K then determines the SNP based on the nucleotide combination with the most reads having matched base at position i. In case of a tie, T1K will select the combination with the least number of SNPs. If the tie could not be broken by the SNP number rule, T1K will report all the equivalent results and mark them as ambiguous. Suppose there are N alleles found for position i, the enumeration step will consider 4^N^ combinations. Since N is small in practice, the enumeration step is still very fast.

### Single-cell processing

T1K supports scRNA-seq data such as SMART-seq data. Since cells could have cell-specific allele expression patterns, we conducted the genotyping for each cell individually and then merge the results. The merge step filters alleles with too low total quality scores and selects the two alleles with the highest quality scores for each gene. After obtaining the allele calling consensus, T1K will remove irrelevant alleles from the reference sequences, and reconduct the genotyping on the reduced reference sequences.

### Allele annotation on HPRC phased genomes

To identify the KIR alleles on a phased reference genome from HPRC in our benchmark study, we align the KIR genomic (exon and intron) sequences from the IPD-KIR to the genome. The first step is to conduct the alignment using minimap2 [41] for quick KIR region discovery. Each KIR region must have at least 99% of bases covered in the alignment interval. Secondly, we realign the sequences from IPD-KIR to each KIR region with BWA-MEM to obtain accurate base-level alignments. Finally, the alleles with the least number of differences in the exonic sequences are selected.

## Abbreviations

MHC: major histocompatibility complex
HLA: human leukocyte antigen
KIR: killer immunoglobulin-like receptor
NK cell: natural killer cell
IPD: Immuno Polymorphism Database
WES: whole-exome sequencing
WGS: whole-genome sequencing
SNP: single nucleotide polymorphism
HPRC: human pangenome reference consortium
1kGP: 1000 Genome Project
LRC: leukocyte receptor complex

## Availability of data and materials

The sample IDs from HPRC and 1kGP used in the study are listed in Table S1 and Table S2.

The SMART-seq data is available at NCBI SRA: PRJNA453180 and PRJNA453183.

T1K source code is available at https://github.com/mourisl/T1K.

The evaluation code for this work is available at https://github.com/mourisl/T1K_manuscript_evaluation.

## Competing interests

X.S.L conducted the work while being on the faculty at DFCI, and is currently a board member and CEO of GV20 Therapeutics. H.L. is a consultant of Integrated DNA Technologies and on the Scientific Advisory Boards of Sentieon and Innozeen.

## Funding

This work is supported by NCI 1R01CA245318 (B.L.), 1R01CA258524 (B.L.), and NIH R01HG010040 (H.L.), U01HG010961 (H.L.), R01HG011139 (H.L.) and U01CA226196 (H.L.).

## Authors’ contributions

L.S, X.S.L, B.L. and H.L. conceived the project. L.S. developed the method. L.S., G.B., B.L and H. L. conducted the evaluations and data analysis. All the authors agree on the manuscript.

## Acknowledgments

We thank the colleagues from X.S.L lab and H.L. lab for helpful discussions.

---

**Table S1.**
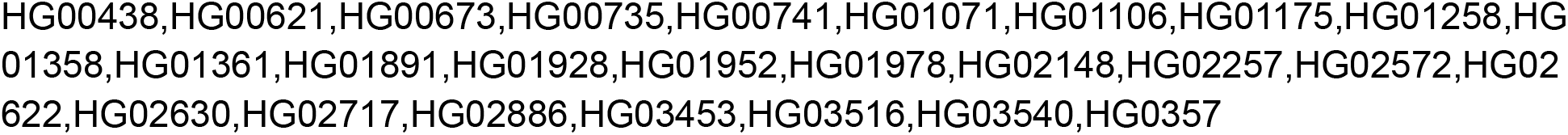
HPRC samples used in the KIR allele prediction evaluation

**Table S2.**
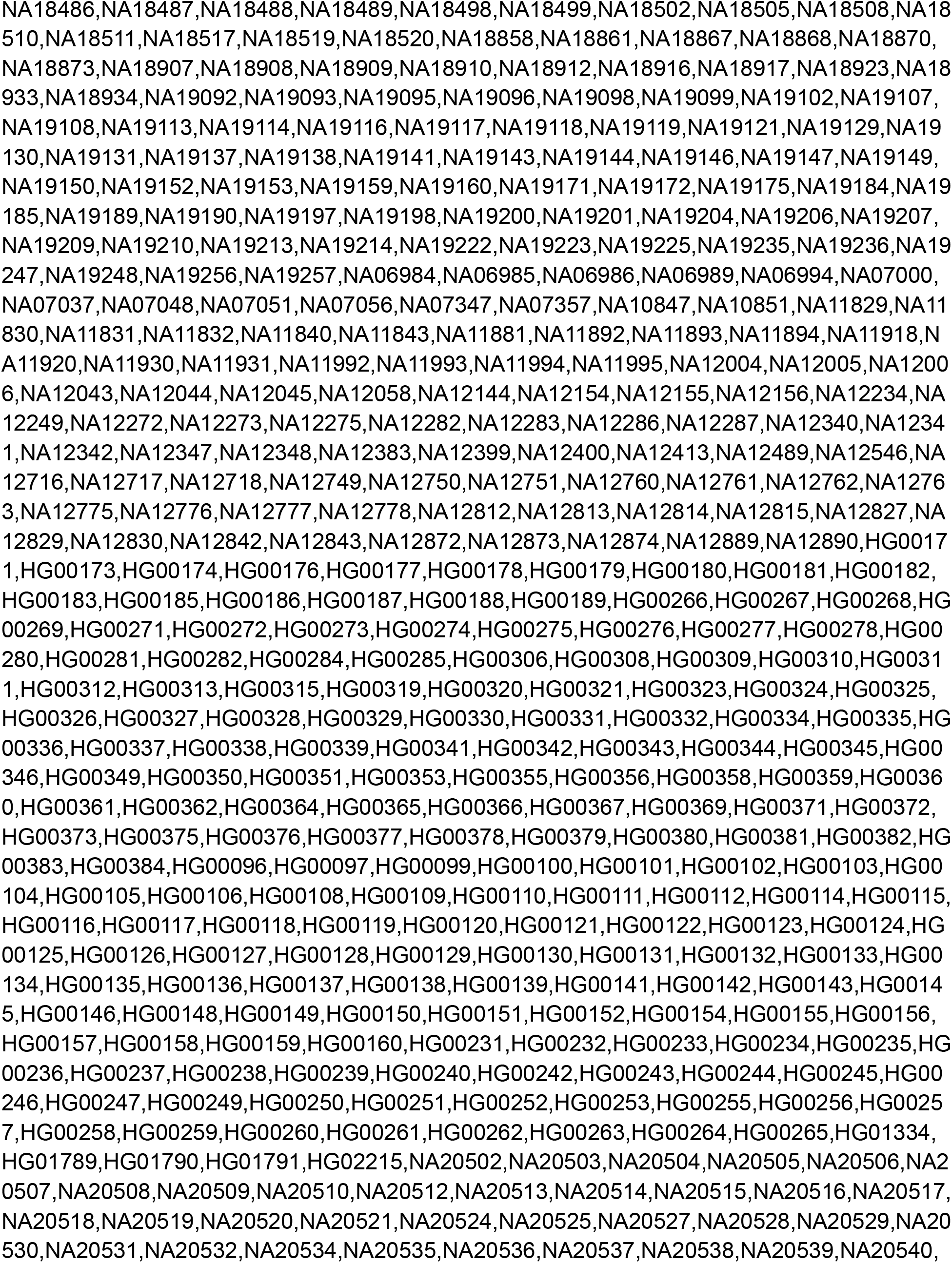
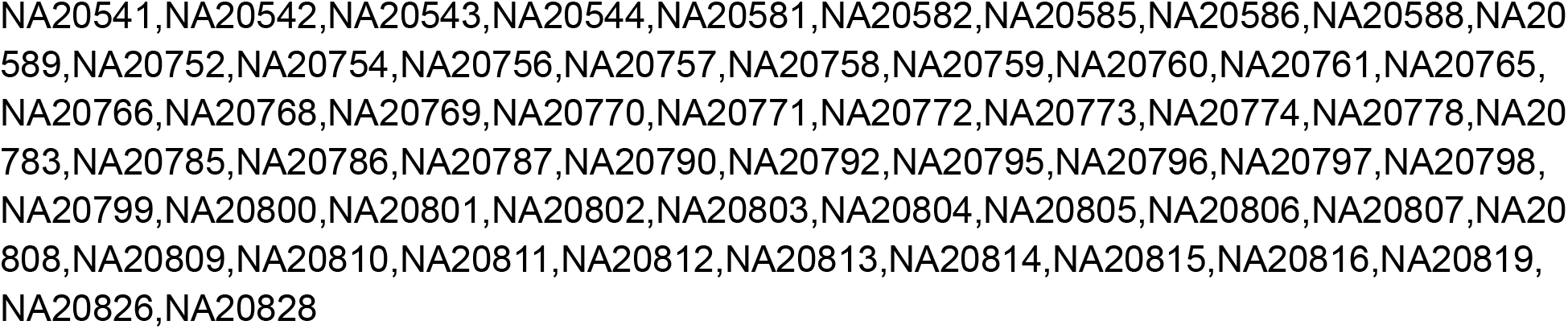
1000 Genome Project samples used in the HLA allele prediction evaluation

**Table S3.** Running time and memory usage of methods on HLA genotyping. All the methods are tested with 8 threads.

|  |  | T1K |  |  | arcasHLA |  |  | OptiType |  |
| --- | --- | --- | --- | --- | --- | --- | --- | --- | --- |
| Sample | # of read pairs (million) | Time (min) | Genotyping time (min) | Mem (GB) | Time (min) | Genotyping time (min) | Mem (GB) | Time (min) | Mem (GB) |
| HG00117 | 246 | 89 | 28 | 36 | 41 | 33 | 21 | 157 | 256 |
| HG00355 | 234 | 78 | 23 | 29 | 29 | 22 | 13 | 103 | 185 |
| NA20527 | 231 | 78 | 22 | 30 | 29 | 23 | 13 | 132 | 219 |
| NA19095 | 228 | 79 | 25 | 29 | 27 | 20 | 12 | 116 | 189 |
| NA06986 | 219 | 85 | 28 | 35 | 31 | 23 | 30 | 175 | 268 |
| NA12716 | 100 | 40 | 15 | 20 | 25 | 23 | 13 | 70 | 137 |
| NA19159 | 98 | 37 | 15 | 18 | 17 | 15 | 16 | 30 | 61 |
| NA12399 | 88 | 31 | 10 | 15 | 18 | 15 | 11 | 39 | 84 |
| NA19108 | 75 | 25 | 9 | 13 | 13 | 11 | 7 | 17 | 38 |
| NA18502 | 71 | 24 | 7 | 10 | 11 | 9 | 6 | 26 | 57 |

**Figure S1:**
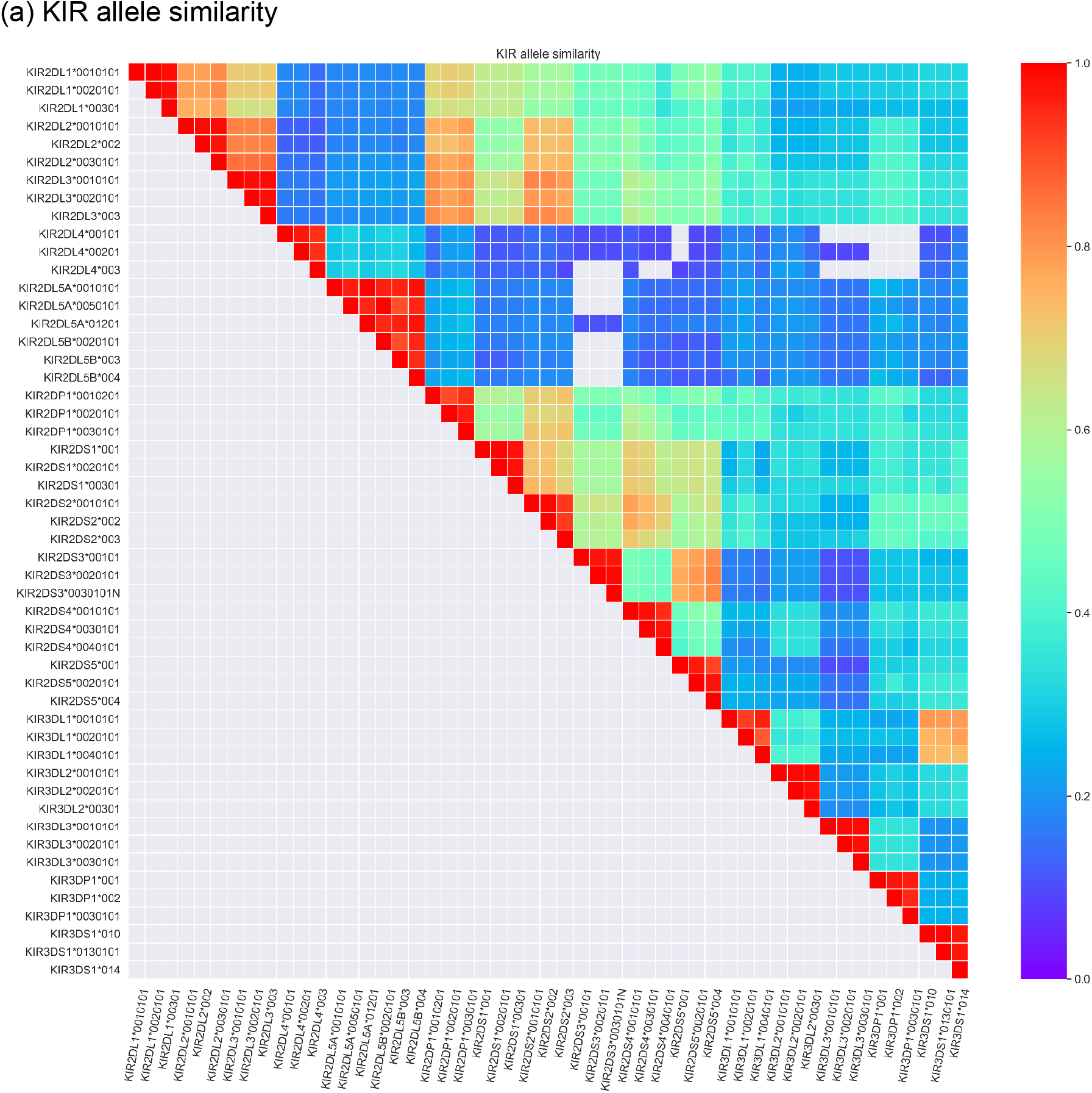
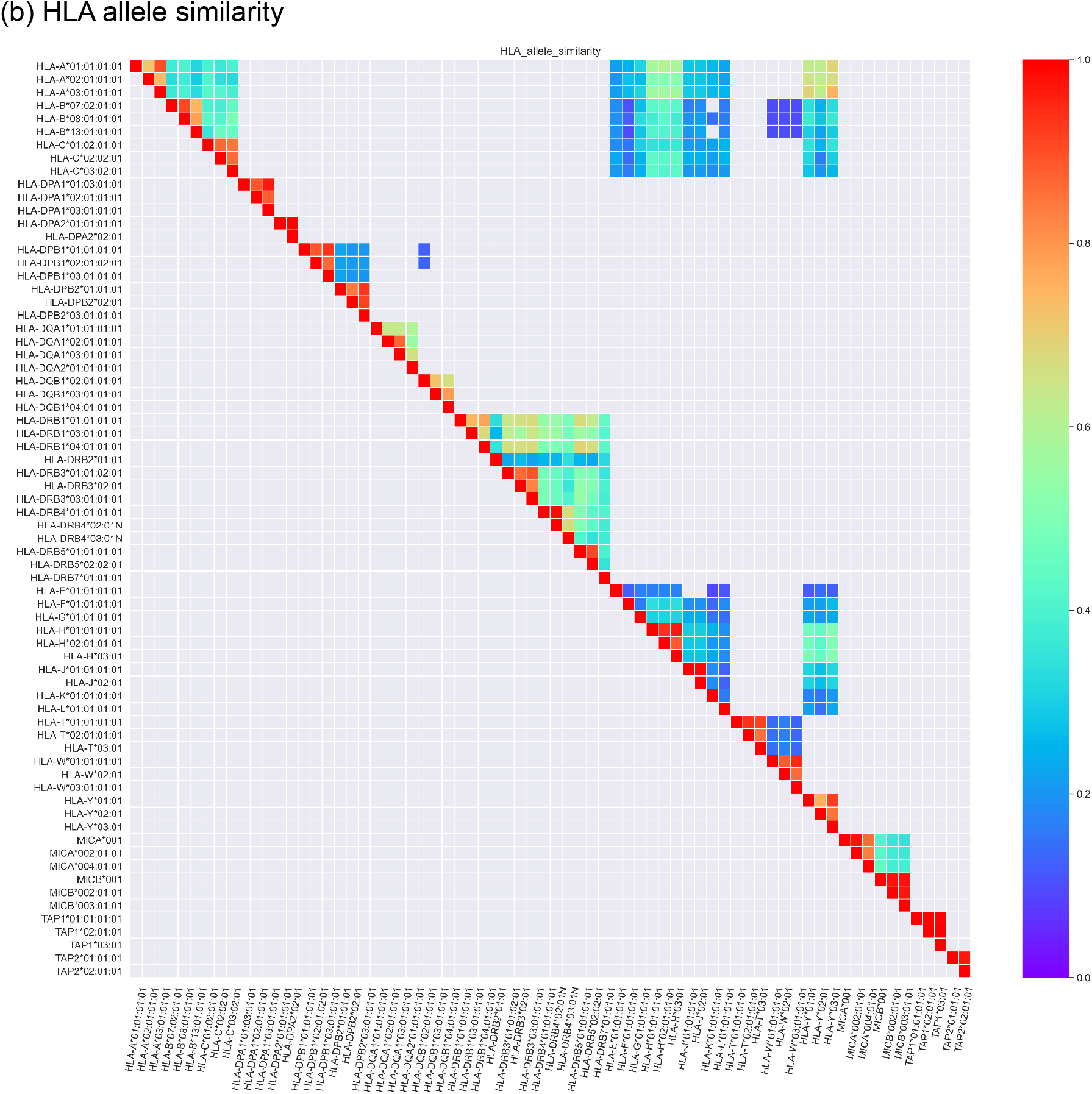
Allele similarity of the mRNA sequences extracted from IPD-KIR and IPD-IMGT/HLA database. Each gene has up to three representative alleles in the heatmap. The color shows the sequence similarity estimated from minimap2 (with -x ava-pb). Only upper triangular portion is colored, and the grey color in upper triangle indicates minimap2 can not find good alignment between two sequences.

**Figure S2.**
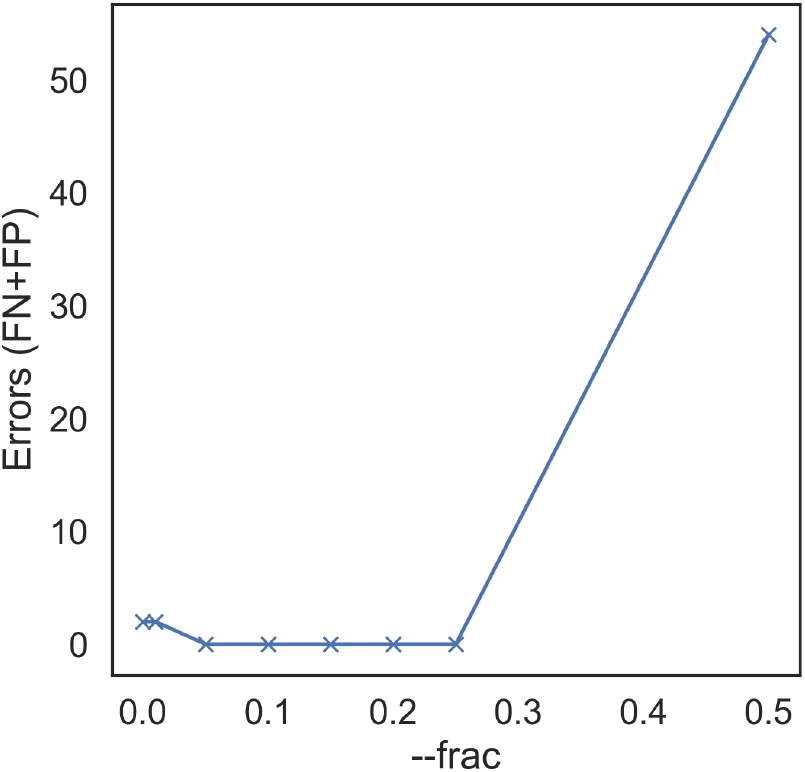
KIR allele calling accuracy when varying the dominant allele fraction threshold (--frac option in T1K) in the simulated data

**Figure S3.**
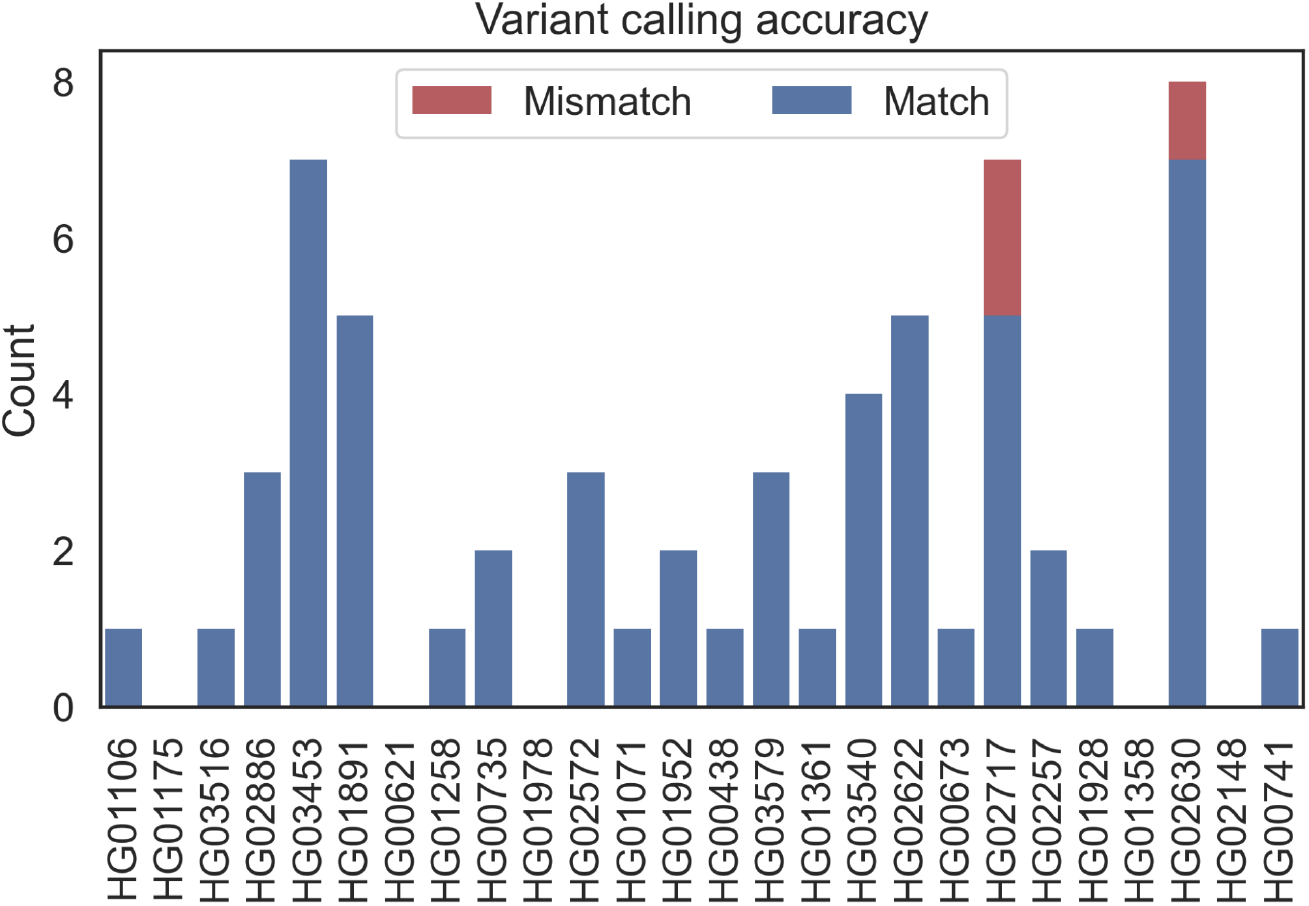
Evaluation of the unambiguous SNPs identified in T1K on the HPRC samples The accuracy is 95.0% (57 validated SNPs over 60 total unambiguous SNPs).

**Figure S4.**
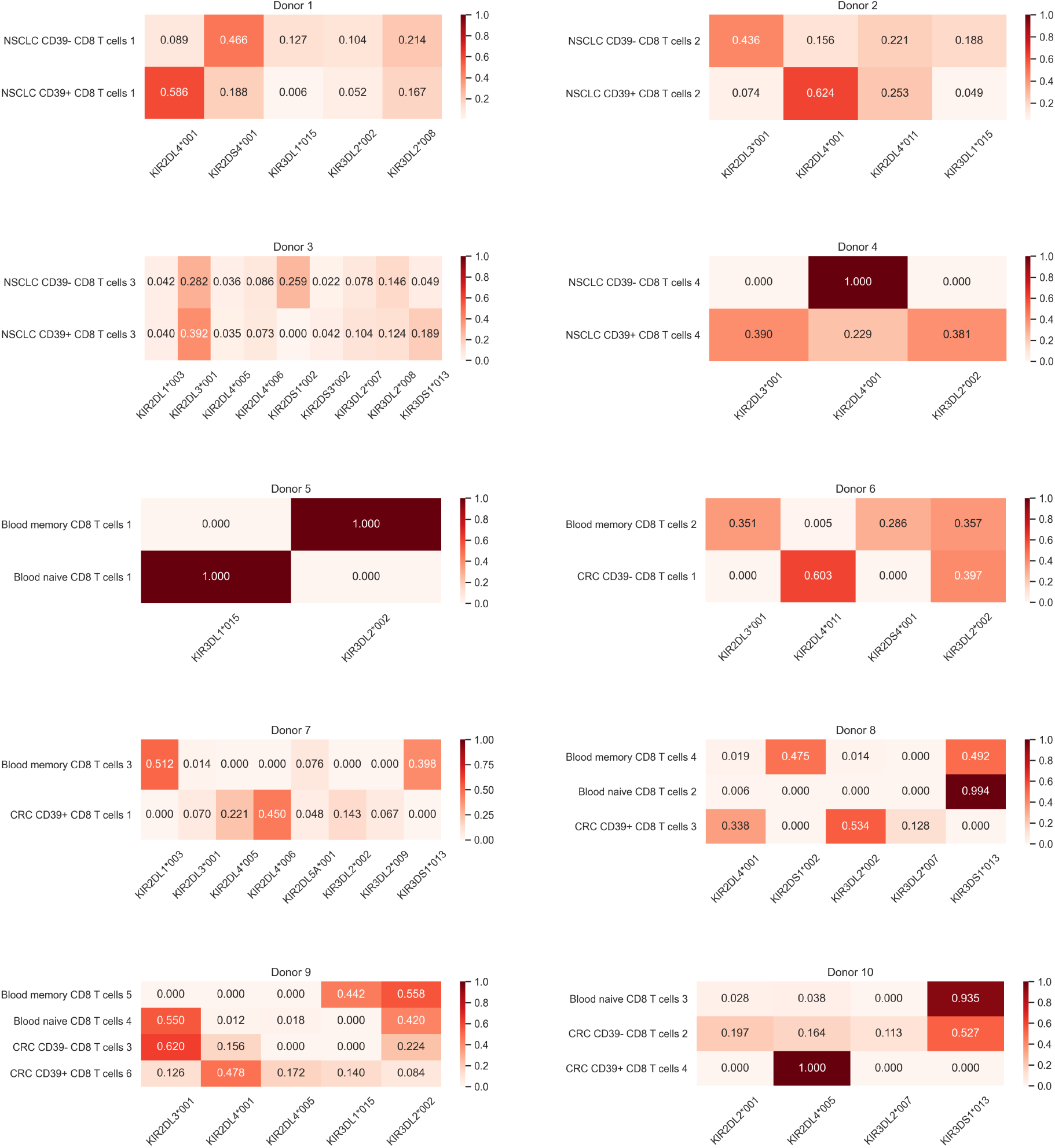
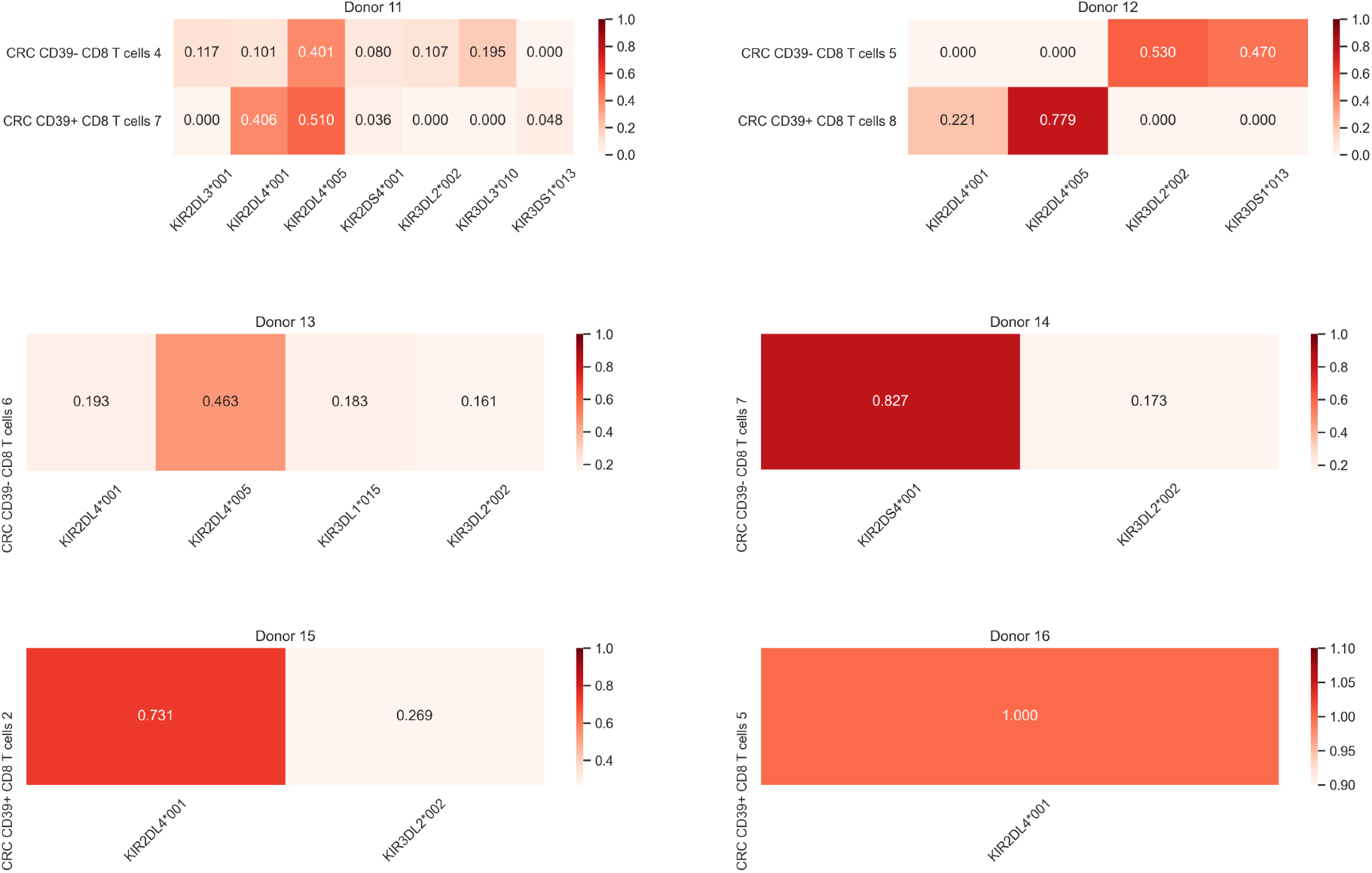
KIR allele expression fractions in SMART-seq scRNA-seq samples on CD8+ T cells. Each heatmap corresponds to the cells bearing the same set of HLA class I genes. Donor 8 is the example in Figure 2a.

**Figure S5.**
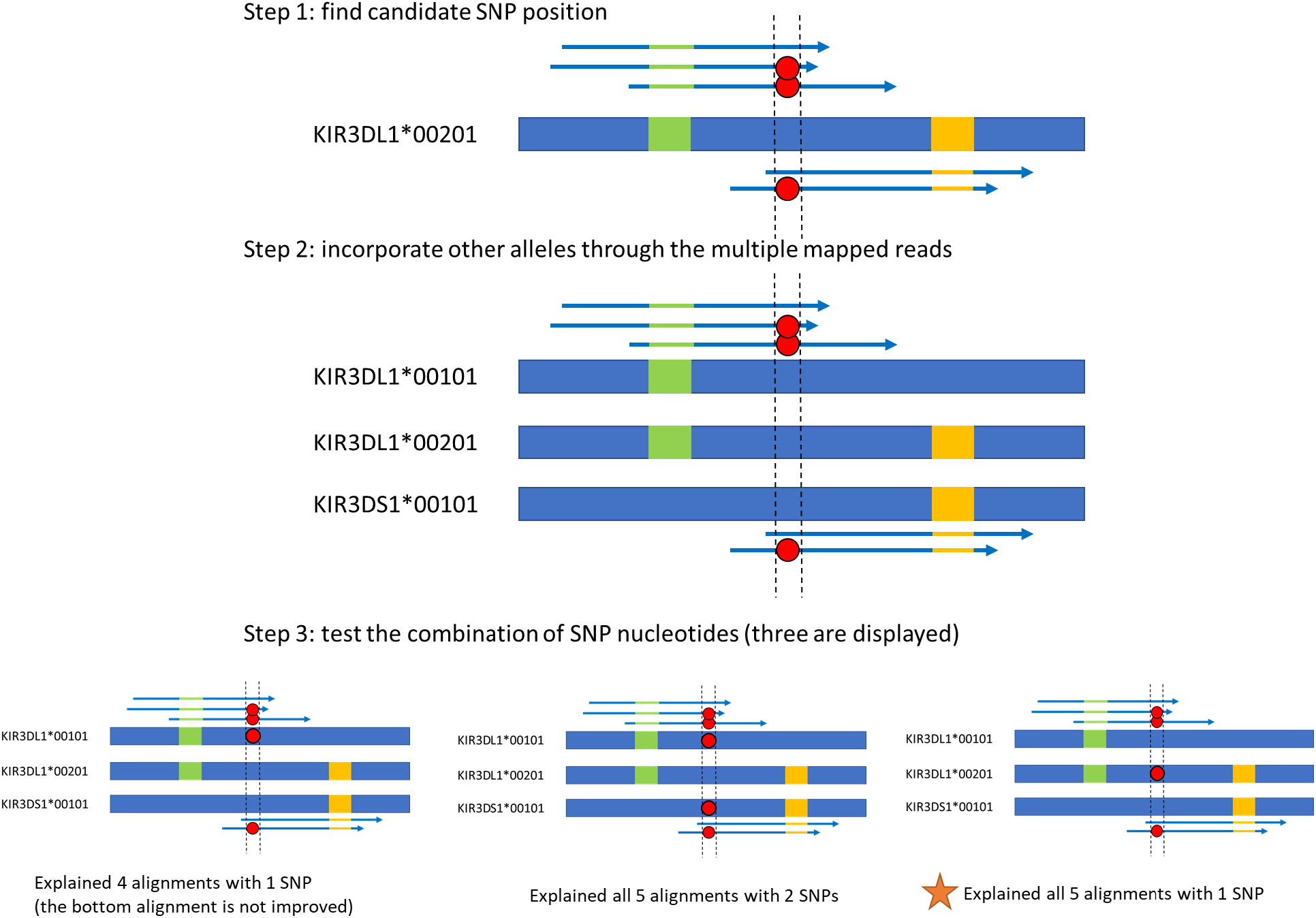
Illustration of the novel SNP identification procedure Green and yellow boxes represent the allele-specific variations. Red dots represent mismatches on the read alignments. The third combination with the orange star is the final SNP result that can improve all five alignments with only one introduced SNP.

